# Neuronal Hyperactivity in Neurons Derived from Individuals with Grey Matter Heterotopia

**DOI:** 10.1101/2024.07.10.602948

**Authors:** Francesco Di Matteo, Rebecca Bonrath, Hanna Schmidt, Ane Cristina Ayo Martin, Veronica Pravata, Rossella Di Giaimo, Danusa Menegaz, Stephan Riesenberg, Femke M.S. de Vrij, Giuseppina Maccarrone, Maria Holzapfel, Tobias Straub, Steven A. Kushner, Stephen P. Robertson, Matthias Eder, Silvia Cappello

## Abstract

Periventricular heterotopia (PH), a common form of grey matter heterotopia associated with developmental delay and drug-resistant seizures, poses a challenge in understanding its neurophysiological basis. Human cerebral organoids (hCOs) derived from patients with causative mutations in *FAT4* or *DCHS1* mimic PH features. However, neuronal activity in these 3D models has not yet been investigated. Here, silicon probe recordings revealed exaggerated spontaneous spike activity in FAT4 and DCHS1 hCOs, suggesting functional changes in neuronal networks. Transcriptome and proteome analyses identified changes in gene ontology terms associated with neuronal morphology and synaptic function. Furthermore, patch-clamp recordings revealed a decreased spike threshold specifically in DCHS1 neurons, likely due to increased somatic voltage-gated sodium channels. Morphological reconstructions and immunostainings revealed greater morphological complexity of PH neurons and synaptic alterations contributing to hyperactivity, with morphological rescue observed in DCHS1 neurons by wild-type *DCHS1* expression. Overall, we provide new comprehensive insights into the cellular changes underlying symptoms of grey matter heterotopia.

## Introduction

Cortical malformations (CMs) result from alterations of one or multiple neurodevelopmental steps, including progenitor proliferation, neuronal migration and differentiation. CMs are frequently associated with epilepsy, cognitive deficits and behavioral alterations^1^, highlighting the importance of better understanding the pathophysiology of these disorders. One of the most common forms of CMs is periventricular heterotopia (PH), which is often associated with seizures and intellectual disabilities and usually is identified by ectopic neurons lining the ventricular zone^2–4^. Genetic causes include *FLNA* mutations and rare variants in several other genes such as *DCHS1, FAT4, ARFGEF2, MAP1B, NEDD4L, ECE2* ^5,6^. Moreover, recently high-throughput analysis of patients with PH indicated an extreme genetic heterogeneity, including copy-number variants^7,8^. As consequences, this heterogeneity might contribute to diversity and variability in disease phenotypes and clinical expression, making the investigation of CMs challenging^9^.

We recently showed that critical features of PH can be modelled in human cerebral organoids (hCOs)^7,10,11^. Causative mutations in the protocadherin-encoding genes *FAT4* and *DCHS1* or knockdown of their expression provoke changes in the morphology of neural progenitor cells (NPCs), contributing to a defective neuronal migration. Neurons were also affected in their migration dynamics and genes involved in axon guidance, synapse organization and ion channel generation were found dysregulated^10^. Interestingly, while knockdown of Fat4 or Dchs1 in mice increases progenitor cell proliferation^12^, this is not the case in PH hCOs, highlighting a species-specific difference^10^.

Inspired by these findings, we here addressed the important question of whether and how these mutations in *FAT4* and *DCHS1* impact neuronal network function. To this aim, we used iPSCs generated from control individuals and two different patients with PH, one was compound heterozygous for mutations in *FAT4* and one homozygous for mutation in *DCHS1* (Supplementary table 6). Complementary studies were performed using isogenic *FAT4* and *DCHS1* KO iPSCs, generated and validated previously^10^, to control for possible differences due to the different genetic background.

## Results

### *FAT4* and *DCHS1* are critical players for the precise formation of neuronal activity

First, we generated 9 months old hCOs from two control individuals, two clinically characterized PH patients with respective *FAT4* or *DCHS1* mutations, and isogenic *FAT4* or *DCHS1* KO cell lines^10,12,13^ (Fig. 1a). We choose this developmental stage as it was shown to consistently exhibit neuronal electrophysiological activity^14^. Immunostainings demonstrated that both FAT4 and DCHS1 hCOs at late developmental stages contain an increased percentage of progenitors (SOX2+) and neurons (DCX+, NEUN+) (Fig. 1b-d), at the expenses of glial cells (NFIA+, S100B+) (Fig. 1e, f) related to control hCOs. The number of both excitatory and inhibitory neurons (EN, IN) was found increased in both PH hCOs (Extended data Fig. 1a-d), however their reciprocal proportion was not altered compared to controls (Extended data Fig. 1e).

**Figure 1.**
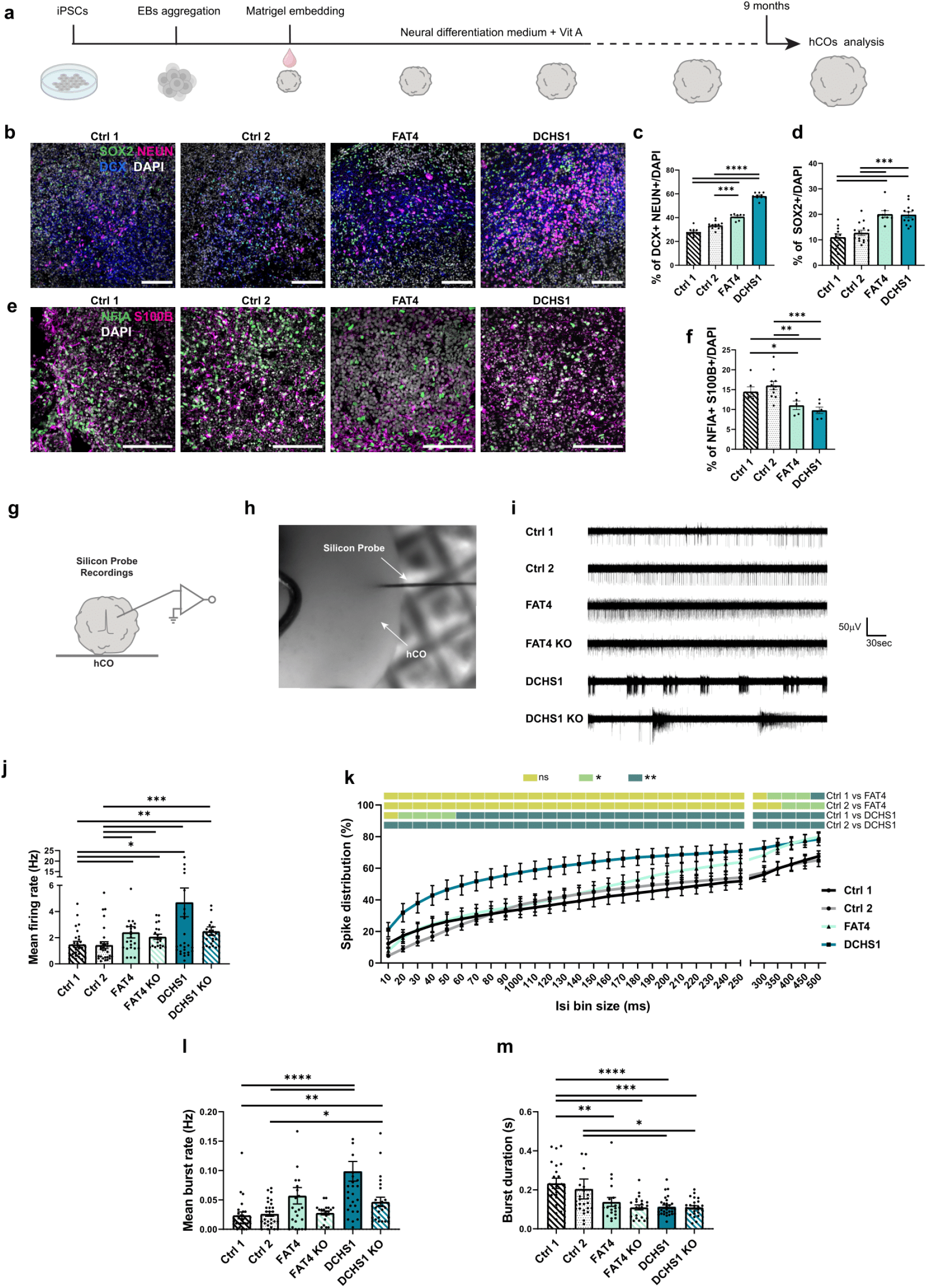
Characterization of control-, patient-derived and KO hCOs. a. Scheme of the generation of hCOs indicating the timepoint used for immunohistochemistry and electrophysiology analysis. Scheme partially created with BioRender.com. b-.Micrographs of 9 months old hCOs sections immunostained for neuronal (DCX, NEUN), progenitor (SOX2) (b) and glial markers (NFIA, S100B) (e) and quantifications of the percentage of positive cells/DAPI (c,d,f) g .Scheme of extracellular silicon probe recordings in an intact hCO. Scheme partially created with BioRender.com. h. Micrographs of a 9 months old hCO during silicon-probe extracellular recording. i .Representative recording traces of spontaneous spike activity recorded in control-, patient-derived and KO hCOs. Recordings were performed for 5 min. j-m. Quantification of the mean firing rate (j), spike distribution (k), mean burst rate (l) and mean burst duration (m) recorded in control-, patient-derived and KO hCOs.Scale bars: 50 µm. Data are represented as mean ± SEM. Statistical significance was based on Mann-Whitney test (*P < 0.05, **P < 0.01, ***P < 0.001,****P < 0.0001). For complete statistical significance refer to (Supplementary table 8). Every dot in the plots refers to independently analyzed recording areas (f) or independent field of view (j).

Next, we performed silicon probe recordings in hCOs (Fig. 1g,h). We developed an experimental approach where hCOs were immobilized in a non-invasive manner, but could be rotated in the recording chamber with a holding frame to allow consecutive insertions of the same probe at different sites of the hCOs (Extended data Fig. 1f). Thereby, we were able to reliably record from spontaneously firing neurons at several locations within a particular hCO. Control experiments confirmed that neuronal firing activity increases upon elevation of the extracellular potassium concentration and vanishes in the presence of tetrodotoxin (TTX) (Extended data Fig. 1g). Intriguingly, both FAT4 and DCHS1 hCOs exhibited exaggerated spontaneous spike activity compared to control hCOs. This neuronal hyperactivity, which was found also in the isogenic FAT4 and DCHS1 KO hCOs, was less pronounced in both FAT4 mutant and FAT4 KO hCOs and was quantified by mean firing rate and inter-spike interval (ISI) analyses (Fig. 1i-k, Extended data Fig. 1h,i). ISI calculations were employed to determine the proportion of high-frequency spikes (defined as consecutive spikes with an ISI <500 ms) in the recordings. In addition, burst analysis revealed an increase of the mean burst rate exclusively for the DCHS1 mutant and DCHS1 KO hCOs, while mean burst duration was found lower in both FAT4 and DCHS1 conditions (Fig. 1l,m). The increase in overall activity and high-frequency spikes suggest that PH hCOs may recapitulate some of the neurophysiological alterations present in PH patients with epilepsy. Intriguingly, these findings suggest also that the mechanisms underlying functional alterations might vary depending on the specific *FAT4* or *DCHS1* mutation.

### Transcriptome and proteome analysis suggest changes in neuronal morphology and synaptic function in PH hCOs

To dissect the two distinct but converging PH phenotypes and to gain deeper insight into molecular changes in FAT4 and DCHS1 neurons, we next performed proteome analysis of control and PH hCOs (Fig. 2a,b, Supplementary table 1). At the protein level, Pearson correlation analysis showed a positive correlation between FAT4 and DCHS1 hCOs related to controls (0.418) (Extended data Fig. 2a), indicating genes dysregulated with the same trend related to controls, such as the synaptic proteins SYT1 and STXBP1 or proteins involved in the neural development such as GAP43 and DPYSL5 that are upregulated in both PH hCOs. However, distinct dysregulated proteins were also detected between the two PH conditions, such as the synaptic proteins SYN1 and SLC1A2 upregulated in DCHS1 hCOs (Extended data Fig. 2b). Gene ontology analysis enrichments for common or specific biological processes and cellular components were identified for FAT4 and DCHS1 hCOs (Fig. 2b,c, Supplementary table 1). For instance, in both PH conditions, fundamental processes like generation of neurons, neuron development and differentiation, axonogenesis, as well as cellular components like synapse and neuron projection were dysregulated. However, FAT4 hCOs additionally showed dysregulated morphological processes, while alterations in the synapse organization was specific to the DCHS1 condition (Fig. 2b,c, Supplementary table 1).

**Figure 2.**
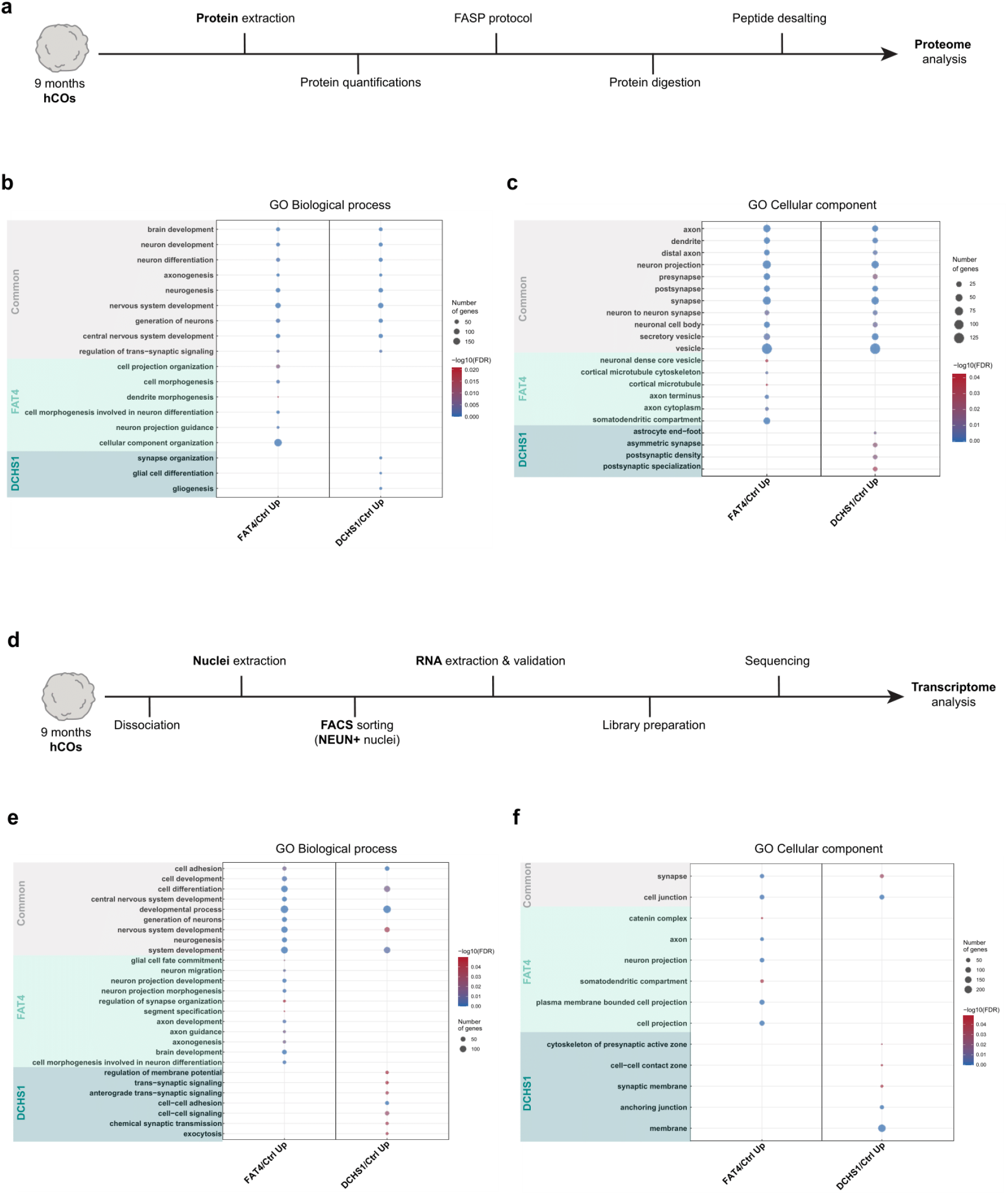
Transcriptome and proteome analysis of control- and patient-derived hCOs. a. Scheme of the samples preparation for proteome analysis. Scheme partially created with BioRender.com b-c. Biological processes (e) and cellular components (f) enriched by GO analysis of the proteome data (Supplementary table 1). Common and FAT4- or DCHS1-specific enrichments are highlighted. d. Scheme of the samples preparation for RNA-Seq analysis. NEUN+ nuclei were isolated from hCOs and processed. Scheme partially created with BioRender.com e-f. Biological processes (b) and cellular components (c) enriched by GO analysis of the RNA-Seq data (Supplementary table 2). Common and FAT4- or DCHS1-specific enrichments are highlighted. Statistical significance was based on Fisher’s Exact.

To better highlight the neuronal alterations suggested by these findings, we next performed RNA-sequencing on NEUN+ nuclei (Fig. 2d). At the transcriptomic level, a positive Pearson correlation was found (0.433) and differences between FAT4 and DCHS1 neurons were identified, such as the voltage-gated sodium channels (VGSCs) genes *SCN3A* and *SCN1A* upregulated in DCHS1 neurons (Extended data Fig. 2c,d). Gene ontology analysis revealed common enrichments in biological processes and cellular components for FAT4 and DCHS1 neurons consistently with our proteome analysis. However, FAT4 or DCHS1-specific enrichments were also similarly detected, highlighting the specific changes at the neuronal level. Indeed, neuron projection morphogenesis, axon development and guidance, somatodendritic compartment were found to be changed for FAT4 neurons, while alterations in the maintenance of intrinsic neuronal properties were specific to the DCHS1 condition (Fig. 2e,f, Supplementary table 2).

Taken together, these changes in molecular signatures provide further evidence for the notion that mutations in different genes, which lead to the same clinical expression, can mechanistically converge and/or diverge at different levels^9^. Moreover, these results let us to speculate that DCHS1 neurons, but not FAT4 neurons, exhibit firing-promoting alterations in intrinsic cell properties, while changes in the spatial cellular organization and interaction, resulting from neuronal morphological and/or synaptic alterations, play a predominant causal role for the FAT4 condition.

### *DCHS1* mutant and KO neurons are more excitable and show a higher density of SCN3A

To test this hypothesis, we first performed patch-clamp recordings in 2D cultures (Fig. 3a).

Control as well as FAT4 and DCHS1 mutant and KO neurons fired solid spike trains upon depolarizing current injections (Fig. 3b, Extended Fig. 3a). Analysis of intrinsic electrophysiological properties revealed no differences between control and FAT4 mutant and KO neurons (Supplementary tables 3 and 4). In contrast, DCHS1 mutant and KO neurons displayed a lower action potential (AP) threshold, a higher AP overshoot and a shorter AP half-width if compared to either control lines (Fig. 3c-f, Supplementary tables 4 and 5). A possible explanation for these findings is that DCHS1 neurons possess a higher density of somatic VGSCs, as suggested from our transcriptome analysis (Extended Fig. 2d). Using immunostainings, we tested this possibility for Na_v_1.3, which is encoded by *SCN3A*. Na_v_1.3 belong to the most highly expressed VGSCs in the human brain and have been associated with epilepsy, intellectual disabilities and CMs^15–17^. Somatic SCN3A signal in DCHS1 mutant and KO neurons was indeed enhanced compared to FAT4 and control neurons (Fig. 3g-h, Extended Fig. 3b). This increase of somatic SCN3A was confirmed also in DCHS1 hCOs (Fig. 3i,j). Using silicon probe recordings, we additionally investigated in DCHS1 hCOs whether the antiepileptic drug and use-dependent VGSC inhibitor lamotrigine^18^ would dampen AP firing in human neurons derived from PH patients (Fig. 3k). Lamotrigine effectively reduced spontaneous spike activity in a dose-dependent manner (Fig. 3l,m).

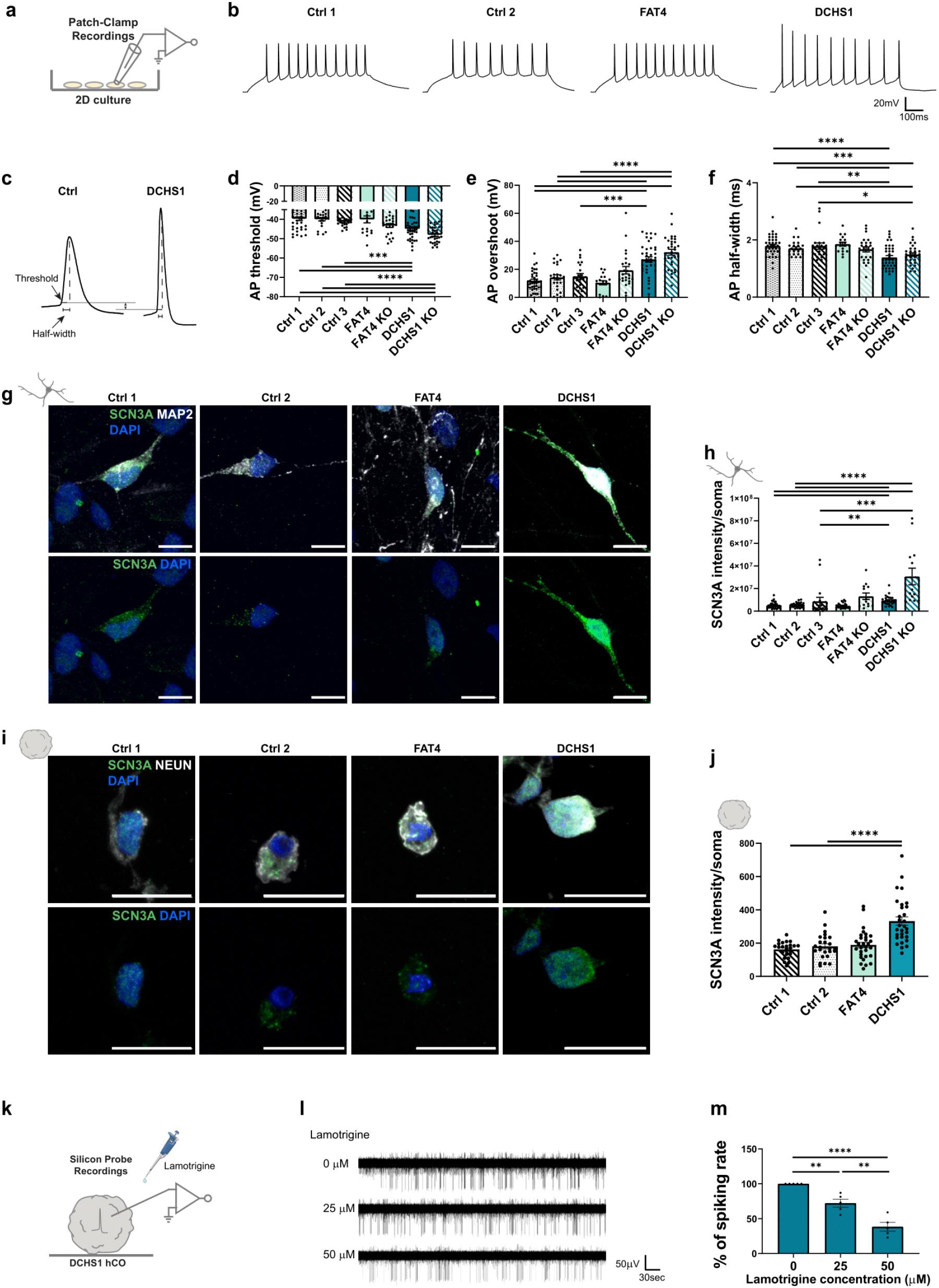
Investigation of single-cell electrophysiological properties and somatic VGSC densities in control-, patient-derived and KO neuronal cultures. a-b. Scheme of whole-cell current-clamp recording in 2D cell culture and representative recording traces depicting evoked neuronal firing. c. Representative APs recorded from a control and a DCHS1 neuron. d-f. Quantification of the AP threshold (d), AP overshoot (e) and AP half-width (f) for control-, patient-derived and KO neurons. g Micrographs of 10 weeks old control- and patient-derived 2D neurons immunostained for NEUN and SCN3A. h Quantification of the somatic SCN3A intensity of control-, patient-derived and KO 2D neurons. I Micrographs of 9 months old control- and patient-derived 3D neurons immunostained for NEUN and SCN3A. j Quantification of the somatic SCN3A intensity of control- and patient-derived 3D neurons. k Scheme of silicon probe recording in a DCHS1 hCO combined with lamotrigine treatment. Scheme partially created with BioRender.com. l Representative traces of neuronal spike activity recorded in a DCHS1 hCO in the absence and presence of lamotrigine. Recordings were performed for 5 min. m Quantification of the percentage of spiking rate in the absence and presence of lamotrigine. Scale bars: 10 µm (g) and 20 µm (i). Data are represented as mean ± SEM. Statistical significance was based on Mann-Whitney test (d-f, h, j) and unpaired ttest (m) (*P < 0.05, **P < 0.01, ***P < 0.001, ****P < 0.0001). For complete statistical significance refer to (Supplementary table 8). Every dot in the plots refers to independently analyzed neurons (d-f, h, j) or independently analyzed recording areas (m).

### *FAT4* and *DCHS1* are key regulators of neuronal morphology and complexity

Next, we infected all four 2D cell cultures (Ctrl 1 and 2, FAT4, DCHS1) with an adeno-associated viral vector (AAV1/2-CMV-eGFP) to sparsely label neurons and, afterwards, performed single-cell morphological reconstructions (Fig. 4a). The morphology of both FAT4 and DCHS1 neurons was altered (Fig. 4b, Extended data Fig. 4a-e) and Sholl analysis demonstrated a higher morphological complexity of these cells (Fig. 4c, Extended data Fig. 4f). Interestingly, the latter phenotype was particularly pronounced for FAT4 neurons (Fig. 4c). We conducted single-cell morphological reconstruction also in 2D isogenic cell cultures (Ctrl 3, FAT4 KO, DCHS1 KO) and confirmed that the altered neuronal morphology and complexity were not the result of different genetic backgrounds (Extended data Fig. 4g-n). Further analysis in 3D hCOs show similar alterations (Fig. 4d-f, Extended data Fig. o-s).

**Figure 4.**
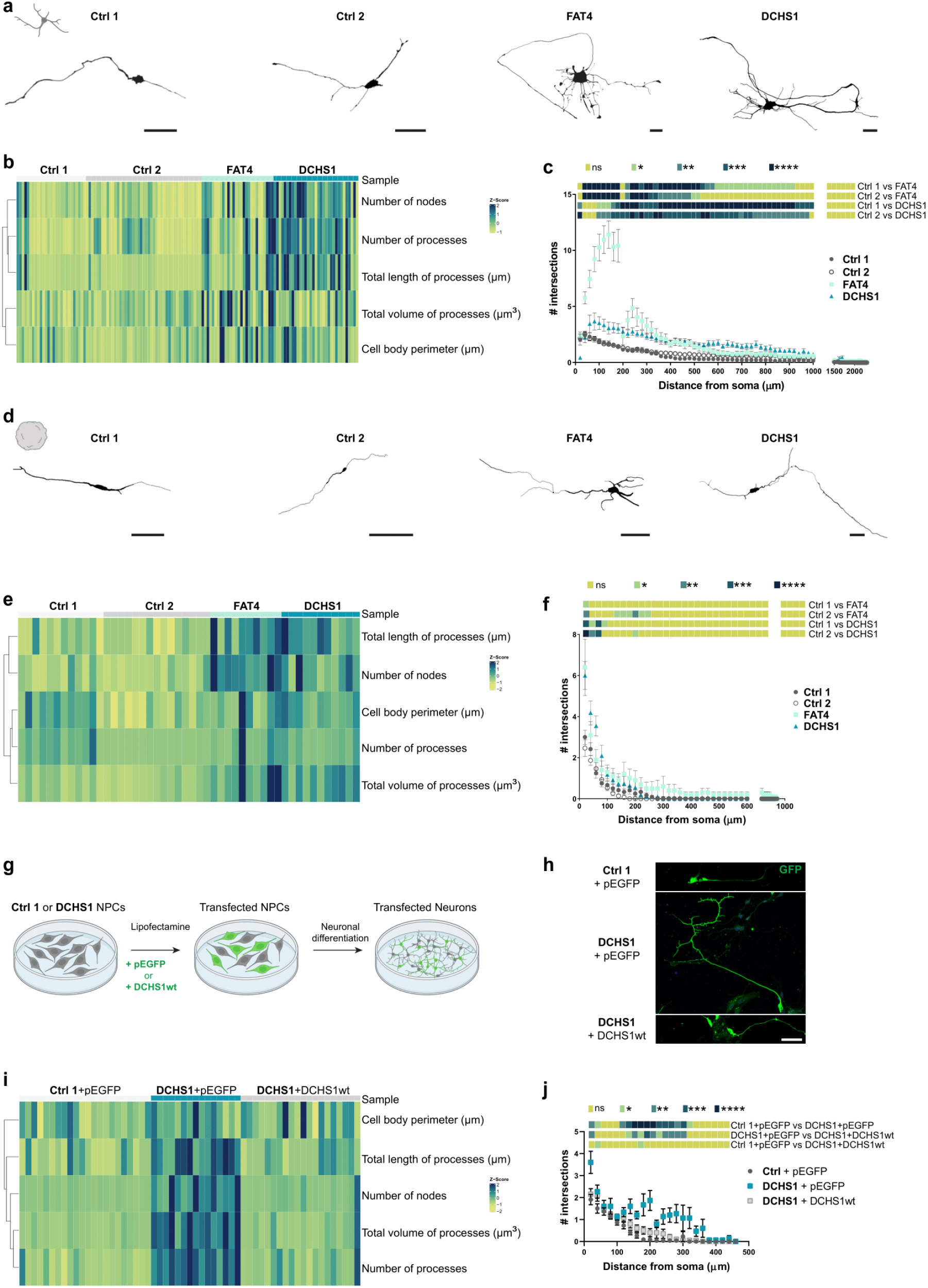
Morphological analysis of control- and patient-derived 2D and 3D neurons. a. Morphologies of representative 10 weeks old control- and patient-derived 2D neurons reconstructed with Neurolucida software. b. Heatmap of the morphological characterization of reconstructed 2D neurons. Z-scores of analyzed parameters are displayed as colors ranging from yellow to blue as shown in the key. c. Quantification of the number of intersections related to the distance from the soma of control- and patient-derived 2D neurons, obtained by Sholl analysis. d. Morphologies of representative 9 months old control- and patient-derived 3D neurons reconstructed with Neurolucida software. e. Heatmap of the morphological characterization of reconstructed 3D neurons. Z-scores of analyzed parameters are displayed as colors ranging from yellow to blue as shown in the key. f. Quantification of the number of intersections related to the distance from the soma of control- and patient-derived 3D neurons, obtained by Sholl analysis. g Scheme of the 2D neuronal differentiation of Ctrl 1 or DCHS1 NPCs upon expression of GFP (pEGFP) or *DCHS1*-wt. Scheme partially created with BioRender.com h. Micrographs of sections of 7 days old control and DCHS1 neurons (9dpt) immunostained for GFP. i. Heatmap of the morphological characterization of control and DCHS1 neurons upon *DCHS1*-wt expression. pEGFP vector was used as control. Z-scores of analyzed parameters are displayed as colors ranging from yellow to blue as shown in the key. j. Quantification of the number of intersections related to the distance from the soma of control and DCHS1 neurons (9dpt), obtained by Sholl analysis. Scale bars: 100 µm (a, d), 50 µm (h). Data are represented as mean ± SEM. Statistical significance was based on Mann-Whitney test (c, f, j) and Fisher’s Exact (b, e, i) (*P < 0.05, **P < 0.01, ***P < 0.001, ****P < 0.0001). For complete statistical significance refer to (Supplementary table 8).

To investigate the role of *FAT4* and *DCHS1* as key regulators of neuronal morphology and complexity, we acutely downregulated *FAT4* or *DCHS1* in control NPCs by transfecting previously validated miRNAs targeting these two genes^10^. Neuronal differentiation was induced 2 days post-transfection (dpt) and morphological reconstructions were conducted on neurons that differentiated for 7 days (Extended data Fig. 4t,u). FAT4 and DCHS1 knockdown neurons displayed changes in their morphology and complexity (Extended data Fig. 4v-ab), demonstrating that *FAT4* and *DCHS1* are modulators of the morphology and complexity of human neurons. Importantly, these morphological alterations were almost entirely rescued by acute overexpression of wild-type (wt) *DCHS1* in DCHS1 neurons (Fig. 4g-j, Extended data Fig. 4ac-ag).

All together, these findings show a key role of *FAT4* and *DCHS1* in the regulation of morphology and complexity of human neurons.

### *FAT4* and *DCHS1* regulate synaptic function

Neuronal morphology plays a prominent role in synaptic connectivity^19^. Our previous single-cell RNA-sequencing analysis performed on FAT4 and DCHS1 hCOs revealed that several genes involved in axon guidance and synapse organization were dysregulated already at early stages of development^10^. Moreover, our proteome analysis showed an upregulation of synaptic proteins in both PH hCOs, including SYN1 which levels were even higher in the DCSH1 condition related to FAT4 (Extended data Fig. 2a,b).

Therefore, we tested for synaptic alterations in 2D cell cultures and 3D hCOs and focused on presynaptic terminals. A significant increase in SYNAPSIN1 and SYNAPSIN2 (SYN1/2) puncta was found in the processes of both FAT4 and DCHS1 mutant and KO neurons (Fig. 5a,b, Extended data Fig. 5a), as well as in both PH hCOs (Fig. 5c,d).

**Figure 5.**
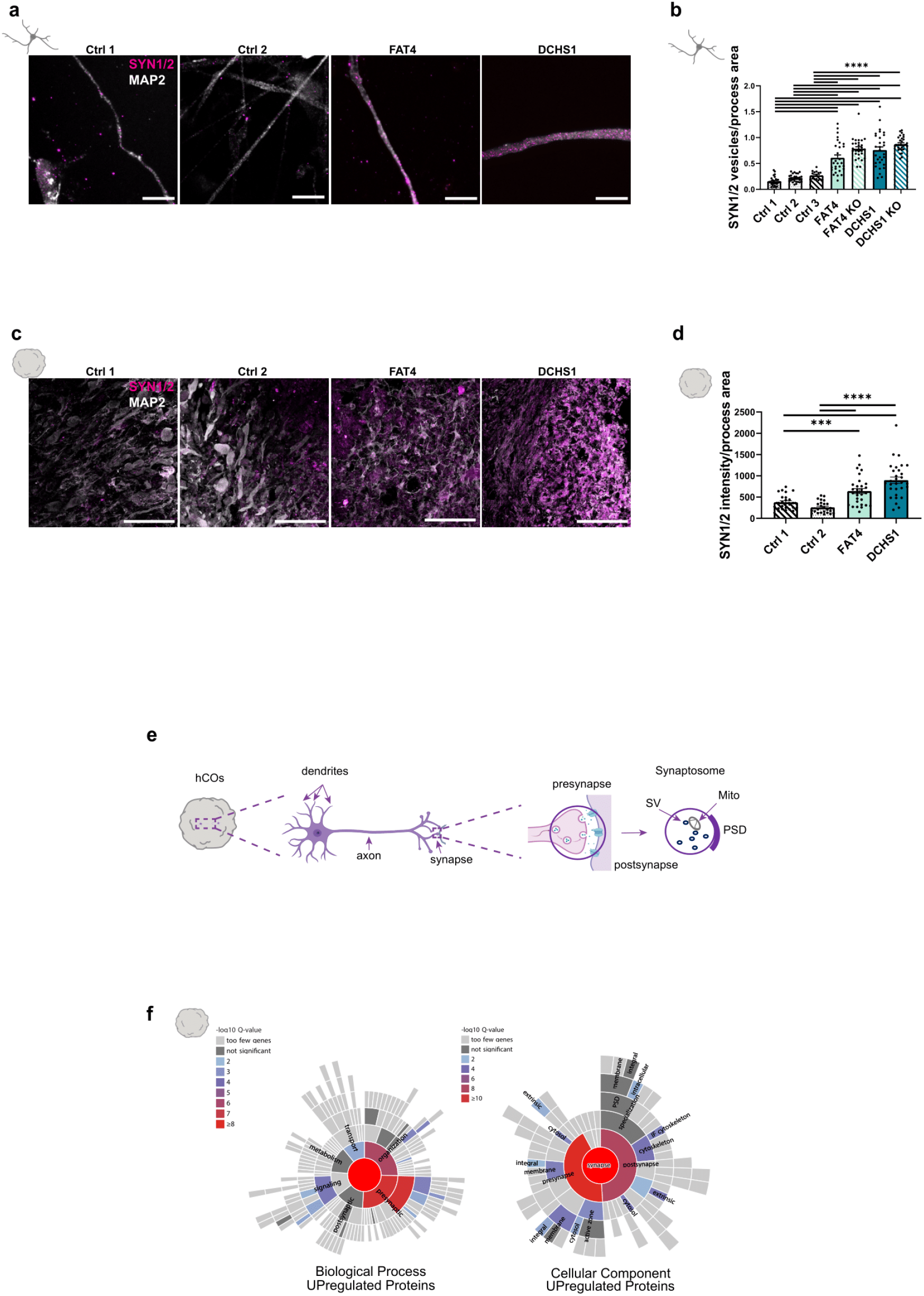
Investigation of synaptic properties in control-, patient-derived and KO neuronal cultures. a. Micrographs of 10 weeks old control- and patient-derived 2D neurons immunostained for MAP2 and SYN1/2. b. Quantification of the number of SYN1/2 puncta related to the MAP2 process area of control-, patient-derived and KO neurons. c. Micrographs of sections of 9 months old control- and patient-derived hCOs immunostained for MAP2 and SYN1/2. d. Quantification of the intensity of SYN1/2 puncta related to the MAP2 process area of control- and patient-derived neurons. e. Scheme of isolation of synaptosomes enriched fractions from hCOs. Scheme partially created with BioRender.com f. Graph showing significantly enriched GO terms of the proteome analysis performed on tissue isolated from hCOs, fractions enriched in synaptosomes. GO analyses show enrichment for biological process and cellular component of proteins upregulated in DCHS1 synaptosomes (Supplementary table 1). Scale bars: 10 µm (a), 100 µm (c). Data are represented as mean ± SEM. Statistical significance was based on Mann-Whitney test (b, d) and Fisher’s Exact (f) (*P < 0.05, ***P < 0.001, ****P < 0.0001). For complete statistical significance refer to (Supplementary table 8). Every dot in the plots refers to independently analyzed neurons.

As exemplarily done for the DCHS1 condition, we also tested for synaptic alterations performing proteome analysis on fractions enriched in synaptosomes obtained from hCOs (Fig. 5e). Synaptosomes represent isolated nerve terminals and are widely used as a model system to study synaptic structure and physiology^20,21^. Most of the dysregulated proteins involved in fundamental biological processes like synapse formation, organization, maturation and neurotransmitter release (e.g., SYN1, SYN3, SYT1) were upregulated in the PH condition (Fig. 5f, Extended data Fig. 5b,c, Supplementary table 1). At the cellular component level, an enrichment of both pre- and postsynaptic compartments was detected (Fig. 5f, Supplementary table 1). Taken together, these findings indicate an involvement of *FAT4* and *DCHS1* in synaptic functions.

## Discussion

In conclusion, we here provide evidences that distinct gene mutations can cause both common and specific cellular alterations that collectively contribute to the clinical expression of PH. This is consistent with the notion that PH result from the dysregulation of different molecular pathways^22^. Although the detailed molecular processes that are controlled by FAT4 and DCHS1 remain to be elucidated, we show that *FAT4* and *DCHS1* mutations, which cause PH, induce an increase of the percentage of both excitatory and inhibitory neurons while their proportions are not altered. We also found that these mutations affect neuronal morphology and complexity and lead to synaptic alterations, indicating a key role of FAT4 and DCHS1 in these processes. These cellular effects, rather than an imbalance of excitatory-inhibitory neuron number, are likely to be causally involved in the neuronal hyperactivity detected in both FAT4 and DCHS1 mutant and KO hCOs, representing a key finding of the present study.

FAT4 and DCHS1 are critical components of the planar-cell polarity (PCP) pathway^23^. PCP proteins play a crucial role in neuronal connectivity by regulating axon guidance, dendritic arborization as well as neuronal migration and polarization^24^. Moreover, neuronal polarization is essential for the asymmetric distribution of proteins between the pre- and postsynaptic compartments^25^. Mutations in several presynaptic and postsynaptic proteins have been linked to epilepsy and small changes in synaptic gain have been demonstrated to lead to seizure-like activity^26,27^.

In contrast, we found alterations in intrinsic electrophysiological properties exclusively for DCHS1 neurons. These changes presumably arise from an increased density of the somatic VGSCs SCN3A and the firing-promoting decrease in AP threshold is likewise compatible with the neuronal hyperactivity seen in hCOs. This finding aligns with the proposal that both gain- and loss-of-function SCN3A mutations may lead to increased seizure susceptibility^15^. Indeed, some pathogenic variants in SCN3A, lead to abnormal brain development and polymicrogyria (PMG) and overexpression of SCN3A in cultured human fetal neurons produced a modest increase in neurite branching without an overall increase in neurite length^16^. Notably, there were also reported infantile epilepsy cases with SCN3A variants not showing PMG^15,28^. These data suggest that SCN3A is important for both, morphological and physiological aspects, during neuronal development. As argued above, this DCHS1-specific phenotype could result from a dysregulation of a distinctive molecular pathway, which is in line with a recent study that identified the previously unknown DCHS1-LIX1-SEPT9 (DLS) protein complex. The DLS complex is crucial to promote filamentous actin organization to direct cell-extracellular matrix alignment and heart valve morphology^29^. Therefore, additional interactors of DCHS1, beyond the well-known interaction with FAT4, might exist.

Overall, we provide detailed new insights into molecular, cellular and physiological alterations that are likely to contribute to the emergence of symptoms in grey matter heterotopia. Moreover, our study supports the use of cerebral organoids for the investigation of human brain development and disorders at different mechanistic levels, aiming for better common or personalized therapeutic interventions.

## Online methods

### Genetic mutation in DCHS1 and FAT4

Details about the genetic mutations found in *FAT4* and *DCHS1* have been previously described^10^ and can be found in the Supplementary Table 6.

### iPSCs culture

Human iPSCs previously reprogrammed from control fibroblasts and peripheral blood mononuclear cells (PBMCs) and from patient fibroblasts (FAT4 and DCHS1) were used to generate neural progenitor cells (NPCs), neurons and hCOs^10,30^. HPS0076 cells were obtained from the RIKEN Bioresource Center, Japan and generated according to the protocol in^31^. KO lines (FAT4 KO and DCHS1 KO) were generated and validate previously from the isogenic control HPS0076 line^10^. CTRL WTC11 iPSC line^32^ was provided by Bruce R. Conklin (The Gladstone Institutes and UCSF) and NPCs were generated from the Steven Kushner lab according to the protocol in^33^. Details about the cell lines used in this study can be found in the Supplementary Table 9.

iPSCs were regularly checked by karyotyping and for mycoplasma regularly and were maintained on Matrigel® Basement Membrane Matrix (Corning, 354234) coated plates (Thermo Fisher), in mTESR1 medium supplemented with 1x mTESR1 supplement (Stem Cell Technologies, 85850) at 37°C, 5% CO2 and ambient oxygen level. The medium change was performed every day. For passaging, cells were treated with Accutase® solution (Life Technologies, A6964) diluted 1:4 in DPBS (Thermo Fisher, 14190144) at 37°C for 5 min. Detached colonies were centrifuged at 300 g for 5 min to collect the pellet and resuspended in mTESR1 with 1x mTESR1 supplement and 10 μM Rock inhibitor Y-27632 (Stem Cell Technologies, 72304) and diluted as needed to the desired density.

### Generation of neurons

Controls and patients NPCs generated previously^10,33^ were differentiated into neuronal cultures following the protocol described in^33^, with small modifications. In short, NPCs were plated on Poly-L-Ornithine (undiluted) (Sigma Aldrich, P4957)/Laminin (50µg/ml) (Sigma Aldrich, L2020) coated 24-well plates (Thermo Fisher) in complete neural differentiation medium consisting in: Neurobasal medium (Life Technologies, 21103049), 1% N2 supplement (Life Technologies, 17502048), 2%B27-RA supplement (Life Technologies, 12587-010), 1% minimum essential medium/non-essential aminoacid (Gibco, 11140-035), 20 ng ml−1brain-derived neurotrophic factor (PeproTech, 450-02), 20 ng ml−1glial cell-derived neurotrophic factor (PeproTech, 450-10), 1μM dibutyryl cyclic adenosine monophosphate (Sigma-Aldrich), 200μM ascorbic acid (Sigma-Aldrich), 2μg ml−1laminin and 1% antibiotic/antimycotic (Sigma-Aldrich, A5955) and cultured for 70 days. The medium change was performed every 2-3 days. For each experiment, many independent neuronal wells (at least 5 wells) from at least three independent batches were analyzed.

### Generation and analysis of hCOs

hCOs were generated as previously described^30,34^. Organoids were kept in 10-cm dishes on an orbital shaker at 37°C, 5% CO2 and ambient oxygen level. The medium changes were performed every 3-4 days. Organoids were analyzed 9 months after the initial plating of the cells. For immunostaining, hCOs were fixed using 4% PFA for 1 h at 4°C, cryopreserved with 30% sucrose, embedded in OCT compound mounting medium for cryotomy (VWR chemicals, 361603E) and stored at -20°C. 16-μm sections of organoids were prepared using a cryotome. For morphological reconstructions, hCOs were embedded in 4% low melting agarose (Sigma-Aldrich, A9414) and 300µm slices were prepared in PBS using a vibratome (speed 0,2mm/s; amplitude 1mm). For each experiment, at least three different organoids for each of three different batches were analyzed.

### Immunohistochemistry

Immunostainings were performed as described previously^30^. Organoids sections and neuronal cell cultures were permeabilized with 0.3% Triton for 5 min and were then blocked with 0.1% Tween, 10% Normal Goat Serum (Biozol, VEC-S-1000). Primary and secondary antibodies were diluted in blocking solution. Nuclei were visualized using 0.5 mg/ml 4,6-diamidino-2-phenylindole (DAPI) (Sigma-Aldrich, D9542). Finally, after several washes with 0.01% Tween in 1X PBS, samples were mounted with Aqua-Poly/Mount (PolyScience, 18606-20) and left to dry in the dark before imaging.

Immunostained sections and neuronal cultures were analyzed using a Leica SP8 confocal laser-scanning microscope with 20X, 25X and 40X objectives. Z-projections were taken to obtain the full 3D image. Notably, for organoids sections a post-fixation step in 4% PFA for 10 min was performed and, for the exposure of nuclei antigen, an extra pretreatment for antigen retrieval before the post-fixation step was performed: The sections were incubated in a freshly made 10 mM citric buffer (pH 6) for 1 min at 720 W and for 10 min at 120 W and then left them to cool down for 20 min at RT. Cells quantifications were performed with the ImageJ software and analyzed with GraphPad. Antibodies list is included in the Supplementary Table 7.

### FACS analysis: nuclei isolation

Nuclei from hCOs were isolated following the protocol from^35^ with small modifications^30^. For each condition, three samples were analyzed as biological replicates; every sample contained three individual hCOs. Briefly, hCOs were dissociated by Dounce homogenization and filtered with BD Falcon tubes with a cell strainer cap (Corning, 352235) to get single cells. RNase inhibitor (NEB, M0314) (0.4 U/ µl) and DNase I (NEB, M0303) (1 U/ µl) were added to the homogenization buffer. RNase inhibitor (0.2 U/ µl) was added to the blocking solution. Primary antibody against NeuN (Millipore, MAB377) was used at 1:1,500 dilution. Secondary antibody Alexa Fluor 546 Goat anti-Mouse IgG1 (c1) (Thermo Scientific, A-21123) was used at 1:2,500 dilution. DAPI (D9542) (0.5 mg/ml) was added during the last 10 min of the secondary antibody incubation. FACS analysis was performed with a FACS Melody TM cell sorter (BD) in BD FACS Flow TM medium, with a nozzle diameter of 100 lm. Debris and aggregated cells were gated out by forward scatter, sideward scatter; single cells were gated out by FSC-W/FSC-A. Gating for fluorophores was done using samples stained with secondary antibody only. Sorted cells were collected in DPBS containing RNAse inhibitor (0.2 units/µl) and 1ml QIAzol® Lysis Reagent (Qiagen, Hilden, Germany) was added to the mix and kept in -80°C until further analysis.

### RNA extraction, cDNA synthesis, and real-time qPCR

RNA extraction, cDNA synthesis and real-time qPCR of NEUN+ and PAX6+ nuclei from FACS sorting were performed as described previously^7^. Briefly, RNA was extracted using RNA Clean & Concentrator Kit (Zymo Research, R1015) and cDNA was synthesized using SuperScript III reverse transcriptase (ThermoFisher, 18080-044) with Random primers (Invitrogen, 48190-11) according to the manufacturer’s protocol. Subsequently, qPCR was performed in triplicates on a LightCycler 480 II (Roche) using the LightCycler 480 SYBR Green I Master (#04707516001, Roche).

The primers sequences were as follows:

GAPDH: FW: 5’-AATCCCATCACCATCTTCCAGGA-3’; RV: 5’-TGGACTCCACGACGTACTCAG-3’

PAX6: FW: 5’-ACCCATTATCCAGATGTGTTTGC-3’; RV: 5’-ATGGTGAAGCTGGGCATAGG-3’^36^

NEUN: FW: 5’-CCAAGCGGCTACACGTCT-3’; RV: 5’-GCTCGGTCAGCATCTGAG-3’^37^

NESTIN: FW: 5’-GGGAAGAGGTGATGGAACCA-3’; RV: 5’-AAGCCCTGAACCCTCTTTGC-3’^37^.

Relative expression was calculated using the DDCp method.

### cDNA amplification library preparation and data analysis

All the steps for the library preparation from the RNA extracted from the NEUN+ were done according to^38^. First, the integrity of the extracted RNA was analyzed at the Bioanalyzer by using the RNA 6000 pico kit (Agilent) following the manufacturer’s protocol.

First-Strand cDNA synthesis and cDNA amplification were done with the SMART-Seq® v4 Ultra® Low Input RNA Kit for Sequencing (Takara Bio USA) following the manufactureŕs protocol. For the cDNA amplification, we performed a range of PCR cycles. The amplified cDNA was purified with AMPure XP beads (Beckman Coulter) following the manufacturer’s protocol. The purified sample was collected and its concentration was measured with the Qubit DNA HS kit following the manufactureŕs protocol. The concentration was ranging between 0.5-2 ng/μl.

The cDNA shearing was performed using the S220 Covaris with AFA technology (Covaris) and the library was prepared with Microplex Library preparation kit v2 (Diagenode) following the manufacturer’s protocol. After the Covaris treatment, the resulting cDNA was in 200–500 bp size range. The concentration, measured with the Qubit DNA HS kit, was 0,5-2 ng/ul (10-20 ng in total). After the library amplification steps with the Microplex Library preparation kit v2 (Diagenode) the DNA concentration was measured again with the Qubit DNA HS Kit. The concentration of the DNA was between 5-6 ng/μl. The library was purified using the AMPure XP beads (Beckman Coulter) and the concentration was checked with Qubit HS DNA kit again, revealing a final concentration between 2-6 ng. Finally, the quality and molarity of the library were also analyzed with the DNA 1000 kit (Agilent). The prepared libraries were sequenced at LAFUGA Genomics, Genzentrum, Universität München (LMU).

### RNAseq analysis

Sequencing reads were aligned to the human reference genome (version GRCH38.100) with STAR (version 2.7.3a). Expression values (TPM) were calculated with RSEM (version 1.3.3). Post-processing was performed in R/bioconductor (version 4.0.3) using default parameters if not indicated otherwise. Differential gene expression analysis was performed with ‘DEseq2’ (version 1.28.1). An adjusted p value (FDR) of less than 0.05 was used to classify significantly changed expression.

### Silicon probe recordings in hCOs

For recordings of spontaneous spike activity in hCOs, a particular hCO was glued with a tiny drop (0.25 µl) of Histoacryl (B.Braun) on a polypropylene mesh that was mounted on a circular plastic frame. Afterwards, the hCO was incubated at 37 ° C for at least 30 minutes in carbogen gas (95% O_2_/5% CO_2_)-saturated ACSF consisting of (in mM): 121 NaCl, 4.2 KCl, 29 NaHCO_3_, 0.45 NaH_2_PO_4_, 20 Glucose, 0.5 Na_2_HPO_4_, 1.1 CaCl_2_ and 1 MgSO_4_. Subsequently, the hCO was transferred with the holding device to the recording chamber and superfused with warm (37°C) carbogenated ACSF at a flow rate of 2.5 ml/min. For additional stabilization, a custom-made anchor was gently placed on the top of the hCO. Recordings were performed using 16-channel probes (Cambridge Neurotech, ASSY-1 E1), which were connected to a ME2100 system (Multichannel Systems). The headstage was mounted on a PatchStar micromanipulator (Scientifica). Recording data were high-pass filtered at 100 Hz, low-pass filtered at 4 kHz, digitized at 20 kHz and transferred via a MCS IFB interface board (Multichannel Systems) to a personal computer. Data were stored using the software Multichannel Experimenter (Multichannel Systems). Visually guided insertion of the silicon probe into hCOs was conducted using a SliceScope microscope (Scientifica) equipped with a 2.5x objective. Once neuronal activity has been detected, the probe was allowed to stabilize in the tissue for 5 minutes and afterwards activity was recorded for 5 minutes. For each condition, independent recording areas were used as biological replicates. Recordings were performed in at least three independent areas per hCO, keeping a distance of at least 400 µm between two adjacent areas. Independent hCOs for each of at least three different batches were analyzed. Sample size: Ctrl 1 n=27; Ctrl 2 n=29; FAT4 n=25; DCHS1 n=29; FAT4 KO n=20; DCHS1 KO n=25. Analysis was conducted with the Offline Sorter^TM^ software (Plexon) and NeuroExplorer software. Burst analysis were conducted following the NeuroExplorer firing rate based algorithm. For experiments with Lamotrigine (Sigma-Aldrich, L3791), the drug was added directly to the ACSF and recordings were started after a waiting period of 5 minutes. For each recording, the channel with the highest activity was used.

### Patch-clamp recordings

Somatic whole-cell current-clamp recordings (>1 GΩ seal resistance, <20 MΩ series resistance, 8 mV liquid junction potential correction, 3 kHz low-pass filter, 15 kHz sampling rate) in iPSCs-derived neuronal cultures were conducted at room temperature (23-25°C) using an EPC9 amplifier (HEKA). Cells were superfused (2-3 ml/min flow rate) with the same carbogenated solution as used for silicon probe recordings. This solution additionally contained NBQX (5 µM) and picrotoxin (100 µM) for current-clamp measurements. Patch pipettes (3-5 MΩ open tip resistance) were filled with a solution consisting of (in mM): 135 KMeSO_4_, 8 NaCl, 0.3 EGTA, 10 HEPES, 2 Mg-ATP, and 0.3 Na-GTP. Current injections were used to depolarize the neuron under investigation. All AP parameters were calculated for the first AP occurring upon depolarizing current injection. Offline analysis was performed using the software FitMaster (HEKA) and Igor Pro (WaveMetrics). AP threshold was defined as the membrane potential where the slope of the AP overshoot reached for the first time a value of 10 mV/ms. For each condition, neurons of independent wells were used as biological replicates. At least three independent wells for each of the three different batches were analyzed. Sample size: Ctrl 1 n=37; Ctrl 2 n=23; Ctrl 3 n=22; FAT4 n=18; DCHS1 n=39; FAT4 KO n=24; DCHS1 KO n=31.

### Morphological reconstruction of neurons and analysis

10-week-old neurons and acute slices of 9-month-old hCOs were used for the morphological analysis. A sparse labeling of cells by using an adeno-associated viral vector (AAV1/2-CMV-eGFP, titer: 2,48×10^12^ gc/mL; produced at the MPIP, transfer plasmid source: Addgene 105530) was performed. 3 days after infection, neuronal cultures and hCOs were fixed at RT for 15 min and 30 min respectively, with 4% PFA and stained for GFP and MAP2 following the described immunostaining protocol. For KD and *DCHS1* expression experiments, 1-week-old neurons were used for the morphological analysis. Neurons were visualized by using a Leica SP8 confocal laser-scanning microscope with 40x objective. Z-projections with a Z-Step Size of 0.50 μm were taken to obtain 3D image. Confocal pictures were used in the Neurolucida Software (MBF Bioscience, Neurolucida version 2017.03.3, 64-bit) and the subsequent tracing was performed in 3D. For the analysis of the reconstructed neurons, Neurolucida Explorer (MBF Bioscience, Neurolucida version 2017.02.9) was used. The “Branched Structure Analysis” tool was used to perform the neuron summary analysis and the individual process analysis. Furthermore, neuronal complexity was obtained using the Sholl analysis tool. To design concentric circles a radius increment of 20 μm were selected. Data were analyzed and visualized on RStudio Complex Heatmap package^39,40^. Statistical tests were performed in GraphPad. For each condition, neurons of independent wells and slices of independent hCOs were analyzed as biological replicates. At least three independent wells for each of the three different batches and three independent hCOs were analyzed. Sample size (2D): Ctrl 1 n=27; Ctrl 2 n=38; Ctrl 3 n=30; FAT4 n=27; DCHS1 n=33; FAT4 KO n=38; DCHS1 KO n=29; Ctrl 1+miRNA NEG n=17; Ctrl 1+miRNA FAT4 n=18; Ctrl 1+miRNA DCHS1 n=14; Ctrl 1+pEGFP n=22; DCHS1+pEGFP n=15; DCHS1+DCHS1 WT n=20. (3D): Ctrl 1 n=12; Ctrl 2 n=15; FAT4 n =10; DCHS1 n =11.

### NPCs transfection

Transfection experiments were performed following the manufacturer’s protocol of Lipofectamine 3000 (Invitrogen, L3000001). Briefly, to knockdown FAT4 and DCHS1, 90% confluent control NPCs were transfected with microRNA against DCHS1 and FAT4 previously generated and validated^10^. To overexpress DCHS1 wt in DCHS1 mutant NPCs, we used the plasmid constructs previously described^10^. DCHS1 wt protein sequence was fused to GFP in pEGFP-C1 plasmid, and the empty vector was used as control. In total, 1 μg of a specific microRNA and plasmid were transfected in NPCs. 2 days post-transfection, NPCs were used to start the neuronal differentiation as described above.

### Quantifications and statistical analyses

Statistical analysis and plotting of data were performed with GraphPad Prism^®^ version 7.04. Statistical significance between unpaired groups was analyzed using *t*-test. Statistical significance between two unpaired groups was analyzed using Mann–Whitney test as indicated in the figure legends. All experiments were reproduced at least three times independently and all attempts at replication were successful. No randomization was performed, but different batches of cerebral organoids for each experiment were used (at least *n* = 3). All acquired data were verified by a second investigator.

### Isolation of synaptosomal enriched fractions from hCOs

Synaptosomal fractions were prepared as previously described^21^ with small changes. Briefly, hCOs were homogenized in nine volumes of cold isotonic medium (HM: 0.32 M sucrose, 10 mM Tris-HCl, pH 7.4), using a Dounce homogenizer. After centrifugation of the homogenate (2,200 g, 1 min, 4°C), the sediment was resuspended in the same volume of HM and centrifuged under the same conditions to yield a washed sediment containing nuclei, cell debris, and other particulates (P1 fraction). The two supernatant fractions were mixed and centrifuged at a higher speed (22,000 g, 4 min, 4°C), to obtain a second sediment that was resuspended in the same volume of HM and centrifuged as described above. The washed sediment contained free mitochondria, synaptosomes, myelin, and microsomal fragments (P2 fraction). The sediment was homogenized in HM and used for subsequent analyses.

### Sample preparation for mass spectrometry

Full lysate and synaptosome enriched samples were resuspended in RIPA buffer, the protein amount was estimated by Bradford protein assay. Protein extract containing 20 ug of protein was subjected to the modified FASP protocol^41^. Briefly, the protein extract was loaded onto the centrifugal filter CO10 kDa (Merck Millipore, UFC201024), and detergent were removed by washing five times with 8M Urea (Merck Millipore) 50mM Tris (Sigma-Aldrich) buffer. Proteins were reduced by adding 5mM dithiothreitol (DTT) (Bio-Rad) at 37°C for 1 hour at dark. To remove the excess of DTT, the protein sample was washed three times with 8M Urea, 50mM Tris. Subsequently protein thiol groups were blocked with 10mM iodoacetamide (Sigma-Aldrich) at RT for 45 min. Before proceeding with the enzymatic digestion, urea was removed by washing the protein suspension three times with 50mM ammonium bicarbonate (Sigma-Aldrich). Proteins were digested first by Lys-C (Promega) at RT for 2 hours, then by trypsin (Premium Grade, MS Approved, SERVA) at RT, overnight, both enzymes were added at an enzyme-protein ratio of 1:50 (w/w). Peptides were recovered by centrifugation followed by two additional washes with 50mM ammonium bicarbonate and 0.5M NaCl (Sigma-Aldrich). The two filtrates were combined, the recovered peptides were lyophilized under vacuum. Dried tryptic peptides were desalted using C18-tips (Thermo Scientific), following the manufacture instructions. Briefly, the peptides dissolved in 2%(v/v) formic acid (Thermo scientific) were loaded onto the C18-tip and washed 10 times with 0.1 % (v/v) formic acid, subsequently the peptides were eluted by 95% (v/v) acetonitrile (Merck Millipore), 0.1% (v/v) formic acid. The desalted peptides were lyophilized under vacuum. The purified peptides were reconstituted in 0.1% (v/v) formic acid for LC-MS/MS analysis.

### Mass spectrometry analysis

Desalted peptides were loaded onto an in-house pulled nano capillary (15 cm, 75 µm ID, Sutter instrument, Puller P-1000), packed with C18 (ReproSil-Pur 1.9 µm, Dr. Maisch GmbH) via the autosampler of the Ultimate 3000R n-LC system (Dionex). Eluting peptides were directly sprayed onto the Q-Exactive Plus Mass Spectrometer coupled to the Nano Spray Flex source (Thermo Fisher). Peptides were loaded in buffer A (0.1% (v/v) formic acid at 300 nl/min and percentage of buffer B (98% acetonitril, 0.1% formic acid) the first 5 minutes solvent B was maintained at 2%, then it was ramped from 2% to 30% over 120 minutes followed by a ramp to 60% over 20 minutes then 98% over 1 minute, and maintained at 98% for another 5 minutes, decreased at 2% over 1 minute and maintained at 2% for 10 minutes. The data acquisition was performed using Xcalibur software (Thermo Fisher). The mass spectrometer was operating in data dependent acquisition mode including one FT survey scan followed by ten HCD MS/MS scans per acquisition cycle. The analysis was performed in the mass range from 375 to 1400 m/z, resolution 70,000 and AGC target 3E6, positive mode. The ionization voltage was 1.9 kV. The HCD fragmentation was performed at resolution 17,500, AGC target 1E5, normalized collision energy (N CE) 27%, max. injection time 100 ms and dynamic exclusion 30s.

### Data Analysis

Raw data were processed using the MaxQuant computational platform (version 1.6.17.0)^42^ with standard settings applied for ITMS Ion trap data. Shortly, the peak list was searched against the Uniprot database of Human database (downloaded in October 2020) with an allowed precursor mass deviation of 10 ppm and an allowed fragment mass deviation of 20 ppm. MaxQuant by default enables individual peptide mass tolerances, which was used in the search. Cysteine carbamidomethylation was set as static modification, and methionine oxidation and N-terminal acetylation as variable modifications. The match-between-run option was enabled, and proteins were quantified across samples using the label-free quantification algorithm in MaxQuant generating label-free quantification (LFQ) intensities.

### Proteomic Analyses

Proteomic data were processed using the RStudio package “DEP”^43^ and following the LFQ-based differential analysis. The MaxQuant output table ‘proteingroups.txt’ was used as input and data were prepared and processed for differential analysis. Result table was then extracted, and results were plotted using RStudio packages ggplot2, dyplyr and tidyverse. An p value of less than 0.05 and a log2FoldChange > 1 for upregulated proteins and <1 for downregulated proteins was used to classify significantly changed expression. For synaptosome fraction analyses, data were further analyzed using SynGO, an interactive knowledge base that accumulates available research about synapse biology using Gene Ontology (GO) annotations to novel ontology terms^40^.

### GO Enrichment analysis

All the GO enrichment analyses were done using Panther^44^ for both proteomics and transcriptomics analyses. We performed a biological process and cellular component overrepresentation analysis for all proteins that were significantly increased (q-value < 0.05), where the corresponding transcript was not significantly increased in expression. Data were then visualized using RStudio script and the ggplot2 R package^45^.

## Data and Code availability

All genomic data will be available at the GEO database under accession number GSE220673.

The mass spectrometry proteomics data will be available at the the ProteomeXchange Consortium via the PRIDE^46^ partner repository with the dataset identifier PXD038760 and 10.6019/PXD038760.

## Acknowledgement

We thank the families participating in this study for their involvement. We thank Barbara Hauger for their excellent electrophysiological technical support; Rosa-Eva Huettl for help providing AAV; Filippo Cernilogar for contribution with transcriptomic data. We also thank all members of the Cappello lab, for technical help, and critical discussions, in particular Marie Lackmann for help with immunostainings and Andrea Forero for help generating schematic drawings. This work was supported by funding from the Max Planck Society, ERA-Net Neuron (nEUrotalk) 01EW1907, ERA-Net E-Rare (HETER-OMICS) 01GM1914, Horizon-EIC-2022-pathfinderopen 3D-BrAIn – 101098791 and by the Netherlands Organ-on-Chip Initiative, an NWO Gravitation project (024.003.001) funded by the Ministry of Education, Culture and Science of the government of the Netherlands and ZonMW PSIDER program TAILORED (10250022110002) (to SAK, FMSDV). Francesco Di Matteo is supported by ERA-Net Neuron (nEUrotalk).

## Author contributions

SC conceived the project. SC and FDM designed the experiments. ME designed the electrophysiological experiments. FDM, RB, HS, AA, DM, and RDG performed the experiments. VP performed proteomic and transcriptomic data analysis. RB performed and quantified immunostainings of hCOs. DM analyzed EPSC recordings. TS helped with the transcriptomic data alignment. FDM, RDG and AA prepared samples for transcriptomic analysis. VP and GM prepared samples for proteomic analysis and validated MS results. SPR provided patients’s fibroblasts. SK and FdV provided control NPCs. MH generated AAV. SR generated the FAT4 and DCHS1 KO lines. FDM, SC and ME wrote the manuscript. All authors provided ongoing critical review of results and commented on the manuscript.

---

**Extended data Figure 1.**
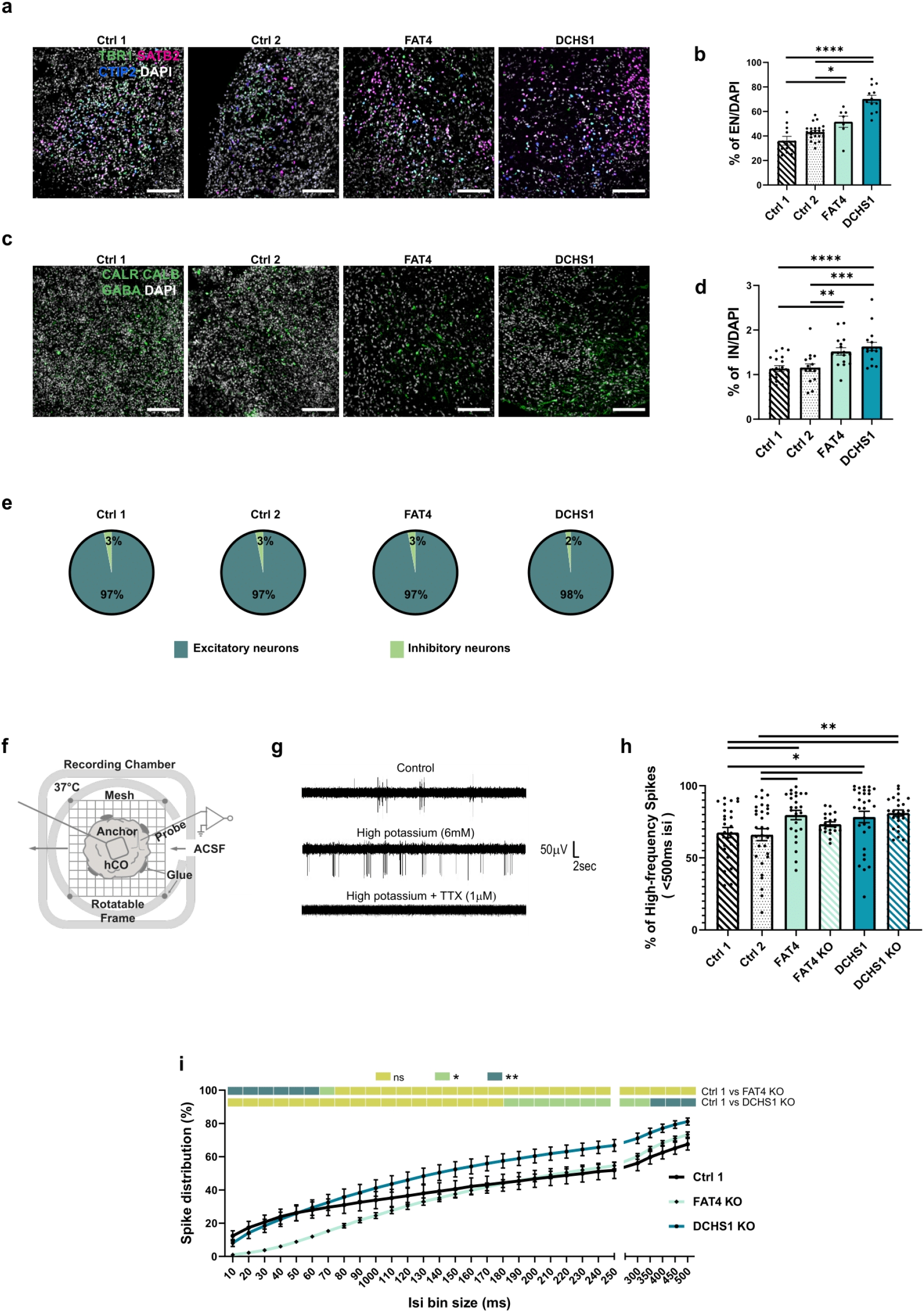
a-b. Micrographs of 9 months old hCOs sections immunostained for excitatory neuron (EN) (TBR1, SATB2, CTIP2) (a) and inhibitory neuron (IN) (CALR, CALB, GABA) markers (c) and quantifications of the percentage of positive cells/DAPI (b,d) e. Quantification of the ratio of EN and IN in control- and patient-derived hCOs. f .Detailed scheme of silicon probe recording in intact hCOs. Scheme partially created with BioRender.com. g .Representative recording traces of spontaneous spike activity in a hCO under different pharmacological conditions (Control, High potassium and High potassium + TTX). Recordings were performed for 30 sec. h. Quantification of the percentage of high-frequency spikes (<500 ms ISI) recorded in control-, patient-derived and KO hCOs. i. Quantification of the spike distribution obtained for control-derived and KO hCOs. Scale bars: 50 µm (a, c). Data are represented as mean ± SEM. Statistical significance was based on Mann-Whitney test (*P < 0.05, **P < 0.01, ***P < 0.001, ****P < 0.0001). For complete statistical significance refer to (Supplementary table 8). Every dot in the plots refers to independent field of view (b) or independently analyzed recording areas (h).

**Extended data Figure 2.**
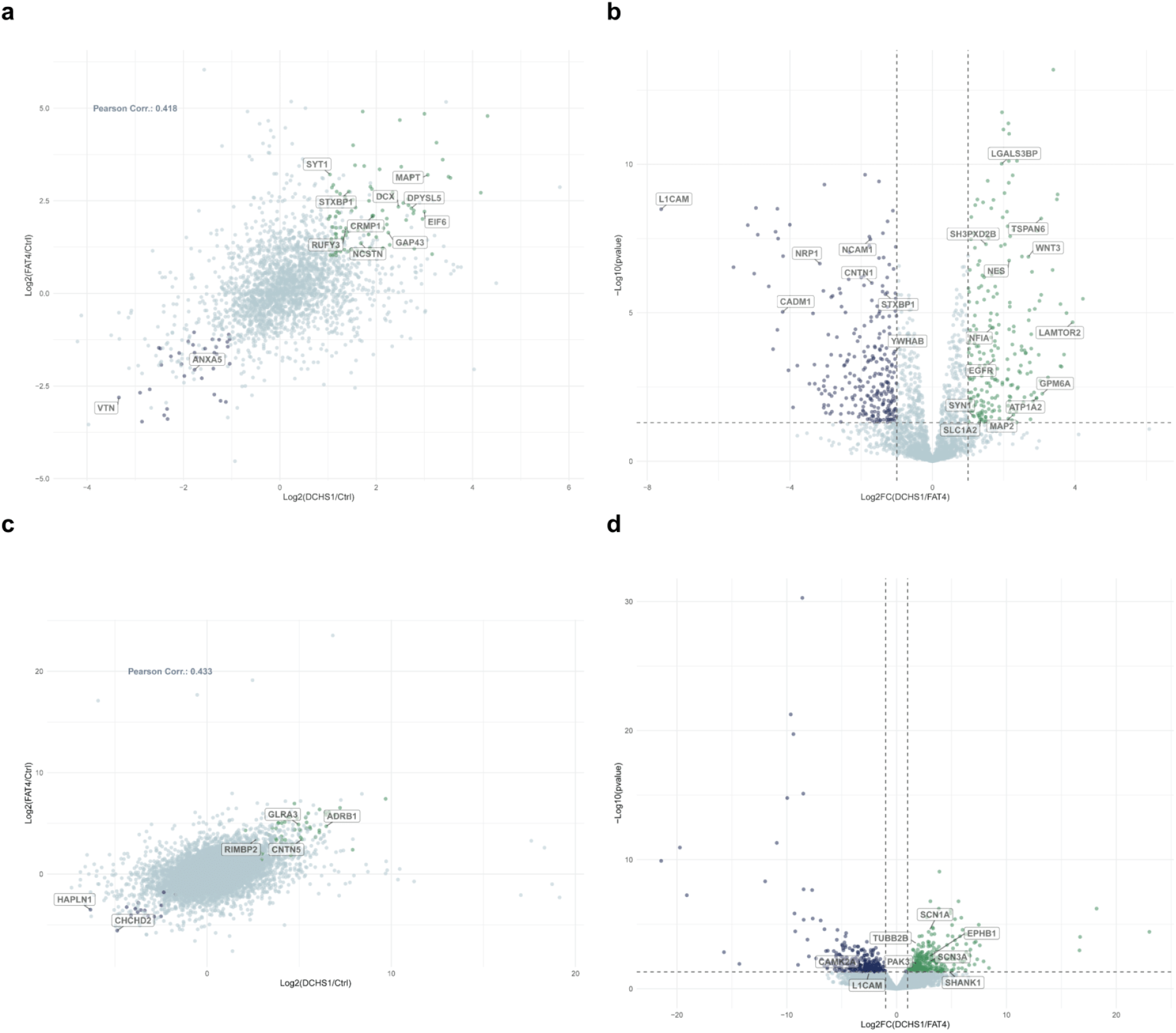
a .Pearson correlation analysis indicating proteins dysregulated with the same trend in the patient’ conditions related to controls. Proteins upregulated are displayed in green, downregulated proteins in blue. b. Volcano plot of dysregulated proteins between FAT4 and DCHS1 hCOs (Supplementary table 1). Proteins upregulated in DCHS1 hCOs related to FAT4 are displayed in green, downregulated proteins in blue. c. Pearson correlation analysis indicating genes dysregulated with the same trend in the patient’ conditions related to controls. Upregulated genes are displayed in green, downregulated genes in blue. d. Volcano plot of dysregulated genes between FAT4 and DCHS1 neurons (Supplementary table 2). Genes upregulated in DCHS1 neurons are displayed in green, downregulated genes in blue. The x-axis represents the log2-fold change in abundance and y-axis the -log10 (p value).

**Extended data Figure 3.**
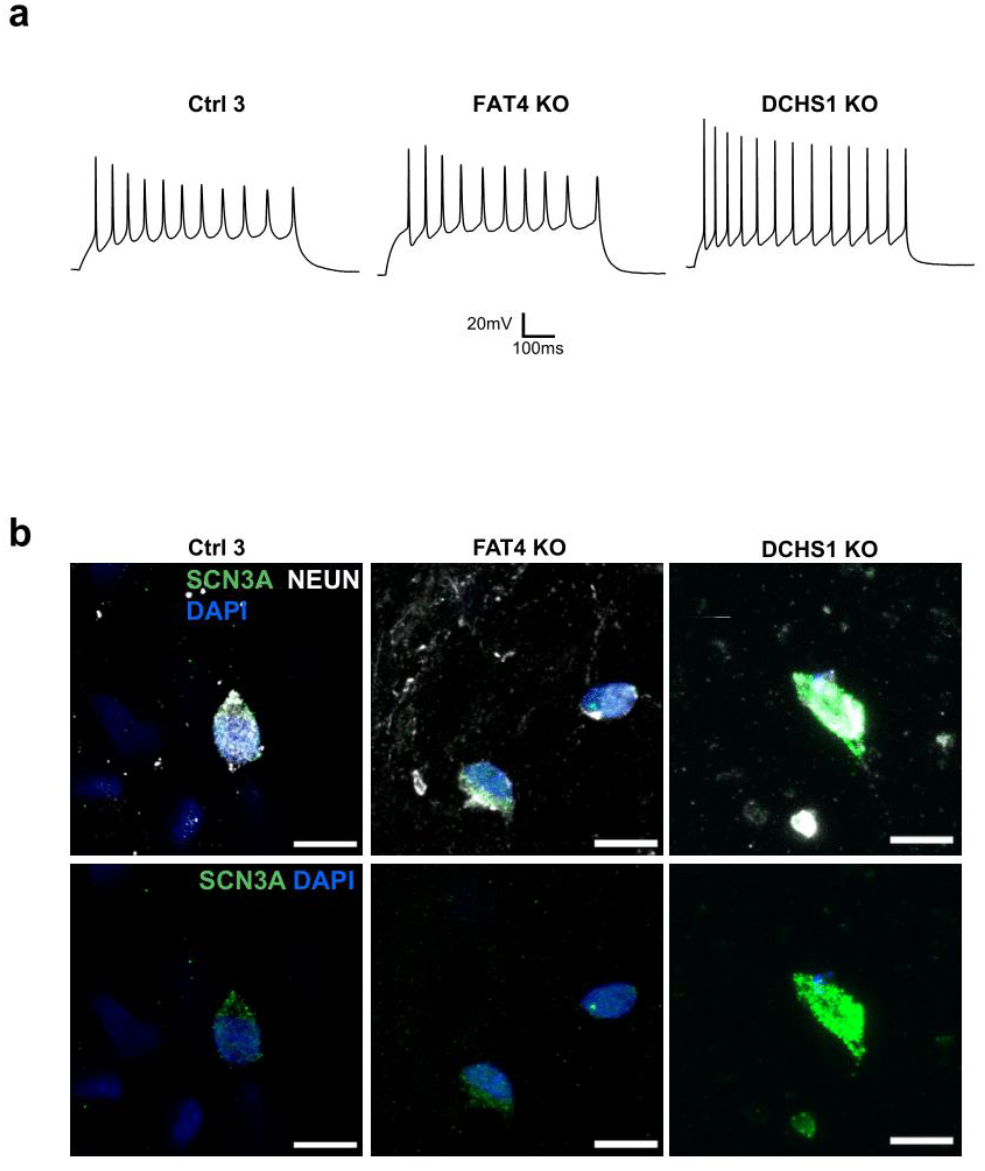
a. Representative patch-clamp recording traces depicting evoked neuronal firing of control-derived and KO neurons. Recordings were performed in current-clamp mode. b .Micrographs of 10 weeks old control-derived and KO 2D neurons immunostained for NEUN and SCN3A. Scale bar: 10 µm.

**Extended data Figure 4.**
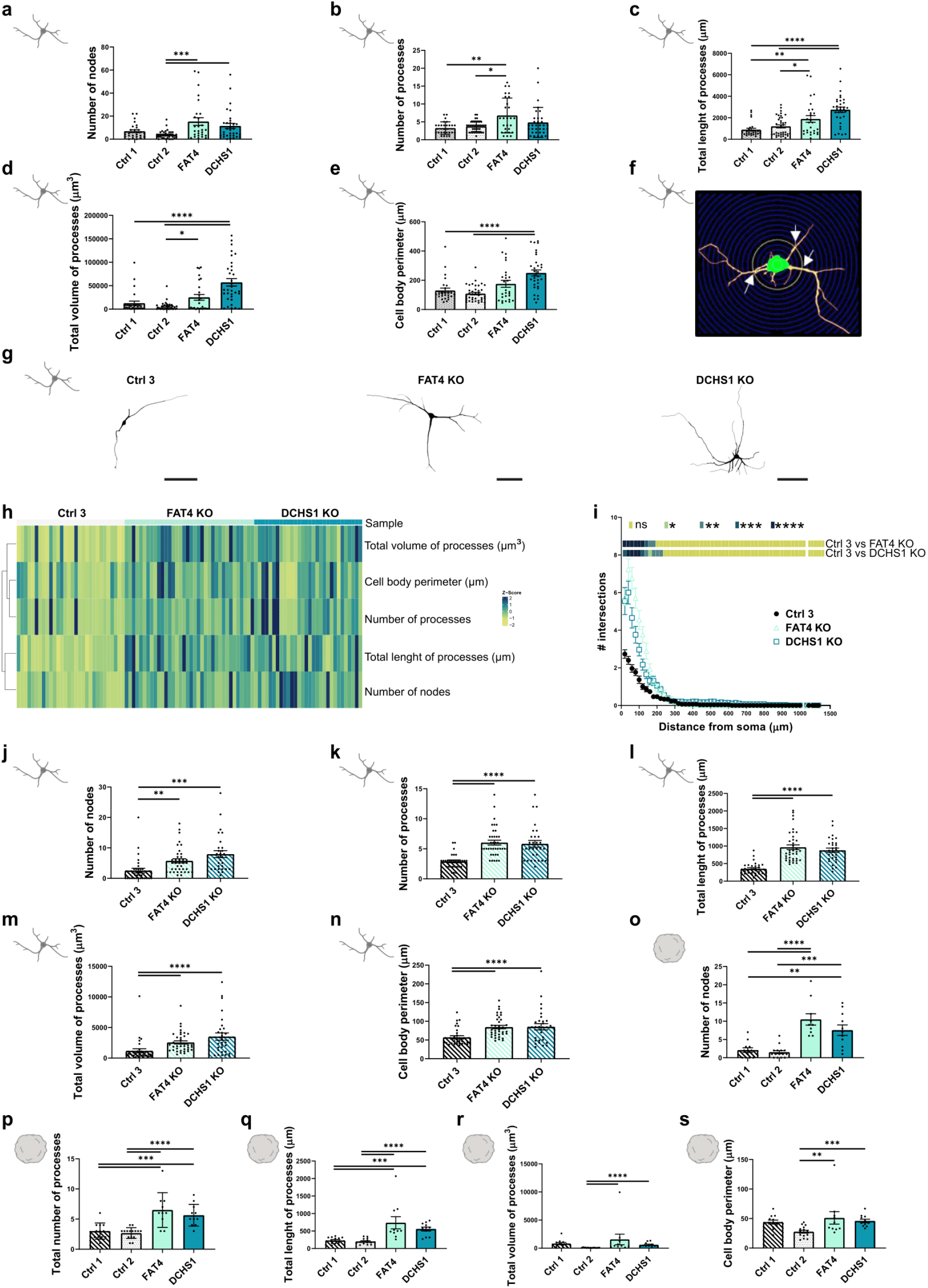
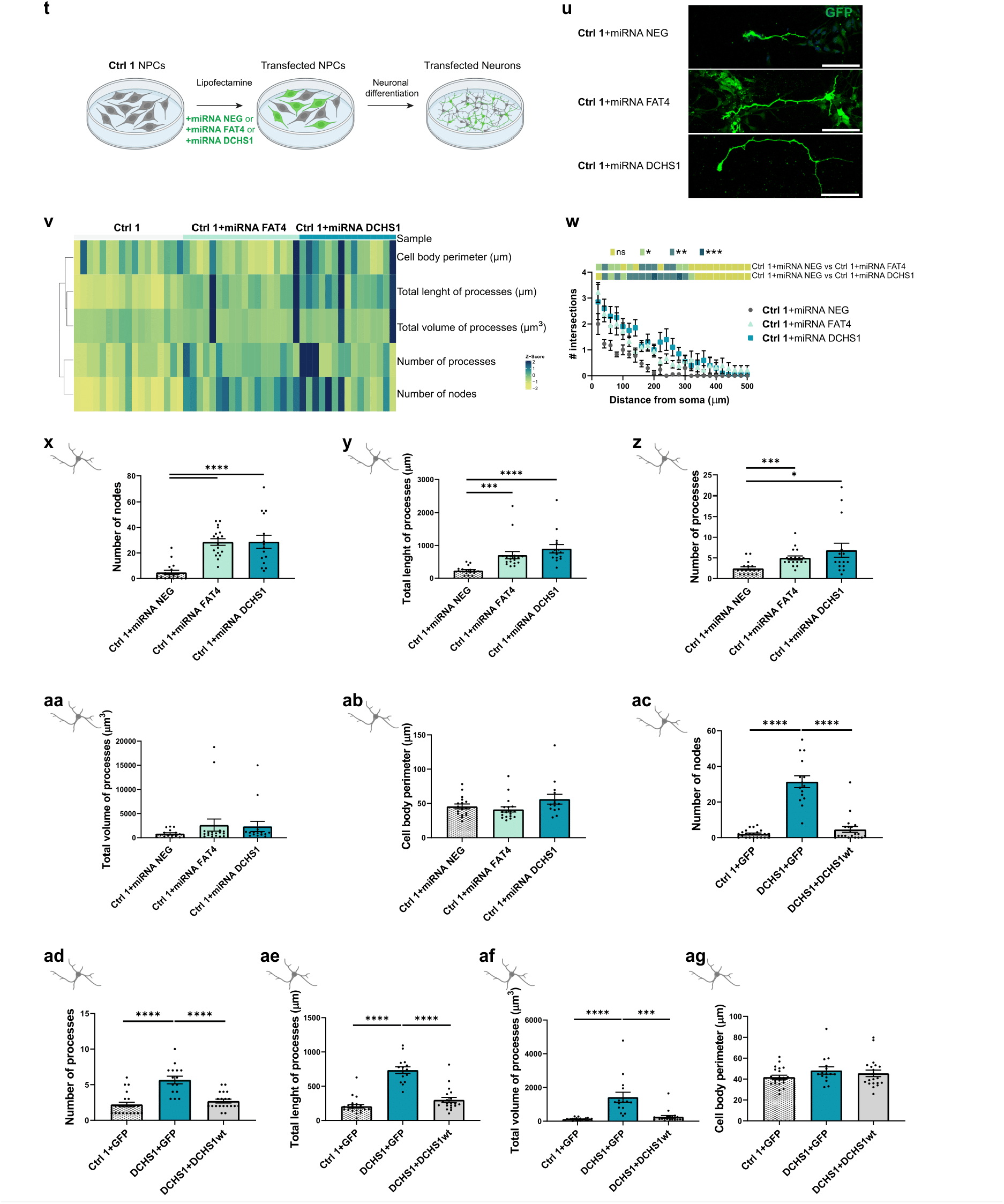
a-e. Quantification of the number of nodes (a), number of processes (b), length of processes (c), volume of processes (d) and cell body perimeter (e) of reconstructed 10 weeks old control- and patient-derived 2D neurons. f. Scheme of Sholl analysis. White arrows show intersections at a certain distance from the soma. g. Morphologies of representative 10 weeks old control-derived and KO 2D neurons reconstructed with Neurolucida software. h. Heatmap of the morphological characterization of reconstructed KO 2D neurons. Z-scores of analyzed parameters are displayed as colors ranging from yellow to blue as shown in the key. i. Quantification of the number of intersections related to the distance from the soma of control-derived and KO 2D neurons, obtained by Sholl analysis. j-n. Quantification of the number of nodes (j), number of processes (k), length of processes (l), volume of processes (m) and c ell body perimeter (n) of reconstructed 10 weeks old control-derived and KO 2D neurons. o-s. Quantification of the number of nodes (o), number of processes (p), length of processes (q), volume of processes (r) and c ell body perimeter (s) of reconstructed 9 months old control- and patient-derived 3D neurons. t .Scheme of the 2D neuronal differentiation of control NPCs upon FAT4 and DCHS1 KD. Scheme partially created with BioRender.com u. Micrographs of 7 days old control neurons (9dpt) immunostained for GFP. v. Heatmap of the morphological characterization of control neurons upon FAT4 and DCHS1 KD. pEGFP vector was used as control. Z-scores of analyzed parameters are displayed as colors ranging from yellow to blue as shown in the key. w .Quantification of the number of intersections related to the distance from the soma of 7 days old control neurons (9dpt) upon FAT4 and DCHS1 KD, obtained by Sholl analysis. x-ab. Quantification of the number of nodes (x), number of processes (y), length of processes (z), volume of processes (aa) and cell body perimeter (ab) of reconstructed 7 days old control neurons (9dpt) upon FAT4 and DCHS1 KD. ac-ag. Quantification of the number of nodes (ac), number of processes (ad), length of processes (ae), volume of processes (af) and cell body perimeter (ag) of reconstructed 7 days old DCHS1 neurons (9dpt) upon *DCHS1*-wt expression. Scale bars: 100 µm. Data are represented as mean ± SEM. Statistical significance was based on Mann-Whitney test (a-e; i-s) and unpaired ttest (w-ag) and on Fisher’s Exact (h, v) (*P < 0.05, **P < 0.01, ***P < 0.001, ****P < 0.0001). For complete statistical significance refer to (Supplementary table 8). Every dot in the plots refers to independent analyzed neurons.

**Extended data Figure 5.**
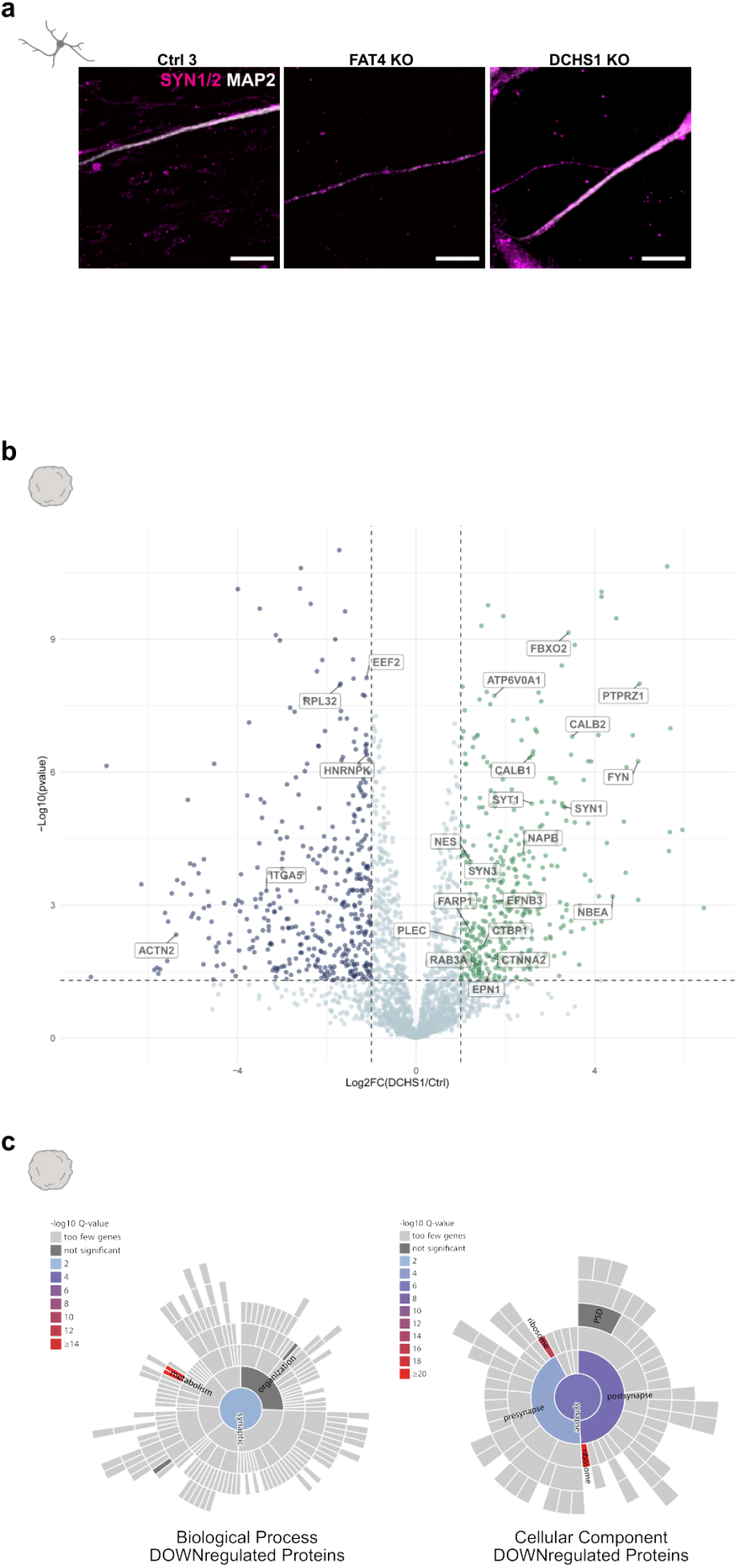
a. Micrographs of 10 weeks old control-derived and KO 2D neurons immunostained for MAP2 and SYN1/2. b. Volcano plot of dysregulated proteins between control and DCHS1 synaptosomes enriched fractions (Supplementary table 1). The x-axis represents the log2-fold change in abundance and y-axis the -log10 (p value). Proteins upregulated in DCHS1 neurons related to control are displayed in green, the downregulated in blue. c .Graph showing significantly-enriched GO terms of the proteome analysis performed on fractions enriched in synaptosomes, isolated from hCOs. GO analysis shows enrichment for biological processes and cellular components of proteins downregulated in DCHS1 synaptosomes (Supplementary table 1). Scale bars: 10 µm. Data are represented as mean ± SEM. Statistical significance was based on Fisher’s Exact.

